# *Arabidopsis PPC2* is crucial for growth at low CO_2_ by involvement in photorespiratory metabolism and integration of ABI5

**DOI:** 10.1101/764589

**Authors:** Lei You, Jumei Zhang, Long Li, Chuanlei Xiao, Xinhua Feng, Shaoping Chen, Liang Guo, Honghong Hu

## Abstract

Phosphoenolpyruvate carboxylase (PEPC) is a pivotal enzyme that plays a key role in photosynthetic CO_2_ fixation in C_4_. However, the function of C_3_ PEPCs and their roles at environmental CO_2_ changes are still limited. Here, we report the role of *PPC2* in seedling growth at low CO_2_ by linking photorespiratory metabolism with primary metabolism and involvement of ABA and ABI5. Mutation of *PPC2* caused seedling growth arrest, with reduced *F*_*v*_/*F*_*m*_, photosynthetic carbohydrates and ABA biosynthesis at low CO_2_. *PPC2* is induced by low CO_2_ and the PEPC activity was greatly reduced in *ppc2* leaves. Moreover, metabolic analyses showed the photorespiratory intermediates, glycine and serine, were greatly increased and primary metabolites were reduced. Application of sucrose, malate and ABA greatly rescued the growth arrest phenotype of *ppc2*. The expression of glycine/serine synthesis and metabolism related photorespiratory enzyme genes were decreased in *ppc2* and regulated by *ABI5* at low CO_2_ conditions. *ppc2* and *abi5* mature plants exhibited reduced A-C_i_ curves at relatively low CO_2_, which could be recovered by non-photorespiratory low oxygen conditions. *ABI5* expression greatly rescued the growth arrest and A-C_i_ curves of *ppc2* at low CO_2_. Our findings demonstrate the important role of C_3_ PEPCs in carbon fixation and metabolism.

## Introduction

Phosphoenolpyruvate carboxylase (PEPC, EC 4.1.1.31) is a ubiquitous enzyme widespread in plants, algae and bacteria (Chollet *et al.*, 1996). There are two functional forms of PEPCs in higher plants, the photosynthetic and non-photosynthetic isoforms. In CAM and C_4_ plants, PEPC plays a pivotal photosynthetic role in primary CO_2_ fixation, by catalyzing the irreversible β-carboxylation of PEP with HCO_3_^−^ to oxaloacetate and inorganic phosphate. The photosynthetic PEPC activity in C_4_ plant has been viewed as essential for carbon assimilation via prefixation of CO_2_ in bundle sheath and, consequently, decreasing photorespiratory activity and resulting in higher water use efficiency (WUE) and higher photosynthetic efficiency than C_3_ plant (Taylor *et al.*, 2018). The non-photosynthetic PEPCs play key roles in plant primary metabolism by replenishing tricarboxylic acid (TCA) cycle to support carbon and nitrogen metabolism (Masumoto *et al.*, 2010), in stomatal opening (Gehlen *et al.*, 1996) and in supplying malate as a substrate of respiration to symbiotic N_2_-fixingbacteroids in legume root nodules (Vidal and Chollet, 1997). PEPC in C_3_ plant is commonly believed to play minor roles in photosynthesis or photorespiration (von Caemmerer, 2013). Recently, NMR analyses of ^13^C-fixation in sunflower indicated that PEPC activity and ^13^C-fixation was significantly increased as net CO_2_ assimilation decreased at high photorespiratory conditions (low CO_2_/O_2_ ratio) (Abadie *et al.*, 2017; Abadie and Tcherkez, 2019). Moreover, mutation of *OsPPC4* in rice led to high accumulation of photorespiratory intermediates, such as glycine, serine and glycerate (Masumoto *et al.*, 2010). Together that photorespiration tightly connects with primary metabolism (Mouillon *et al.*, 1999; Rachmilevitch *et al.*, 2004), these studies give clues that C_3_ PEPC may also play roles in photorespiration or photosynthesis.

In *Arabidopsis*, there are four genes encoding PEPC. Mutation in either *AtPPC1*, *AtPPC2* or *AtPPC3* led to decreased fresh weight and delayed flowering time (Feria *et al.*, 2016). Moreover, mutation of both *AtPPC1* and *AtPPC2* greatly reduced malate and citrate synthesis, and severely suppressed ammonium assimilation, which finally led to the growth arrest of *ppc1ppc2* (Shi *et al.*, 2015). However, how these *AtPPCs* regulates plant primary metabolism and whether these *AtPPCs* regulate photosynthesis and plant development under different stress conditions, such as photorespiratory low CO_2_ conditions, are yet unknown.

The phytohormone abscisic acid (ABA) play a prominent role in the establishment of stress tolerance. In addition, ABA regulates important aspects of plant development, including inhibiting embryo and seed development, promoting seed desiccation tolerance (Yamaguchi-Shinozaki and Shinozaki, 2006), influencing seeds dormancy, germination and seedling establishment (Finkelstein *et al.*, 2002), and facilitating vegetative development. ABA regulates these processes through affecting the expression levels of corresponding genes, which are modulated by various ABA-responsive trans-acting factors, such as B3-domain family proteins (e.g. *ABI3, VAL1*) (Giraudat *et al.*, 1992; Suzuki *et al.*, 2007), APETALA2 (AP2) family proteins (e.g. *ABI4*) (Finkelstein *et al.*, 1998) and basic leucine zipper (bZIP) family proteins (e.g. *ABI5*) (Finkelstein and Lynch, 2000). ABI5 is a key component in ABA-triggered pathways during germination, and seedling establishment, as well as subsequent vegetative growth (Lopez-Molina et al., 2001). It has been reported the role of ABI5 in nitrogen assimilation and signaling. *abi5* mutant seedlings displayed decreased sensitivity to nitrate inhibition (Signora *et al.*, 2001; Yang *et al.*, 2011). ABI5 also have been reported to positively regulate chlorophyll catabolism related genes, *SGR1* and *NYC1*, through recognizing their upstream ABA response elements (ABREs) element (Sakuraba *et al.*, 2014). Therefore, ABI5 is proposed to be a key player in monitoring environmental conditions during seedling growth. However, whether ABI5 is responsive to CO_2_ changes and functions in low CO_2_-induced plant growth are unknown.

In this study, we demonstrate the crucial role of *AtPPC2* in seedling growth at limiting CO_2_ conditions by linking photorespiration metabolism with primary metabolism, via integration of ABI5. *ppc2* mutant showed retarded seedling growth at low CO_2_ conditions and reduced carbon assimilation, in which *ABI5* expression was heavily suppressed. Expression of *ABI5* rescued the growth arrest phenotype of *ppc2* at low CO_2_ and reduced photosynthesis and biomass at ambient CO_2_. Metabolic analyses showed photorespiratory intermediates glycine and serine were accumulated, and malate were decreased in *ppc2* at low CO_2_ conditions. Our study demonstrates *PPC2* and *ABI5* are key regulators in plant growth at limiting CO_2_ conditions in C_3_ plants, which make positive contribution to carbon fixation and metabolism.

## Materials and Methods

### Plant Material and Growth Conditions

All *ppc* mutant lines used in this study were in the Columbia (Col-0) background. The mutant lines *ppc1* (SALK_088836) (Feria *et al.*, 2016), *ppc2* (SALK_128516) (Shi *et al.*, 2015), *ppc3* (SALK_143289) (Feria *et al.*, 2016) were obtained from the Arabidopsis Biological Resource Center (http://abrc.osu.edu) and their homozygous were confirmed by PCR (Table S1).

All seeds were surface-sterilized and sown on 1/2 Murashinge and Skoog medium (MS) (pH 5.7). After cold treatment, seeds were germinated and grew in Percival Chambers with different CO_2_ conditions (200 ppm, 400 ppm, respectively), at a light regime of 16 h light /8 h dark (light intensity 100 μmol m^−2^ s^−1^) and relative humidity (RH) of 56%.

### Generation of Constructs and Transgenic Plants

The coding sequences of *PPC2* and *ABI5* were amplified from *Arabidopsis* cDNA with primers PPC2-OE-F/PPC2-OE-R, and ABI5-OE-F/ABI5-OE-R, respectively. PCR products were cloned into pGreen-35S and pEarlyGate-35S-YFP (Earley *et al.*, 2006). The promoter region of *PPC2* amplified by primers Pro-PPC2-F and Pro-PPC2-R from *Arabidopsis* genome was cloned into the vector pEarlyGate-100-GUS (Earley *et al.*, 2006). All primers used in construct generation were presented in Supplemental Table 1. The constructs were introduced into Arabidopsis by *Agrobacterium tumefaciens*-mediated transformation using the floral dip method.

### Calculation of Chlorosis Rate

15-day-old seedlings of Col-0 and *ppc2* mutant which were growing at 200 ppm and 400 ppm CO_2_ condition were analyzed for chlorosis rate.

Chlorosis rate (%) = number of chlorotic seedlings / number of total seedlings

### PEPC Activity Assays

0.1 g leaves (fresh weight) of 15-day-old seedlings growing at 200 ppm and 400 ppm were harvested. Tissues were ground and extracted in 1ml extraction buffer. After centrifugation, the supernatant was immediately used for PEPC activity detection by a PEPC activity assay kit (Nanjing Jiancheng Bioengineering Institute). 1 unit of PEPC activity was defined as 1 nmol of NADH oxidation per min and per mg protein at 25◻. Total protein content was quantified by using BCA (bicinchoninic acid) protein assay kit (Sangon Biotech).

### A-C_i_ curve analyses

The measurements of A-C_i_ curves were performed by a closed infrared gas exchange analysis system (LI-COR 6400XT). 4 to 5-week-old leaves were clamped in the 2 cm^2^ chamber with leaf temperature at 21°C for measurement. The measurement of A-C_i_ curves was performed at increasing CO_2_ concentrations of 50, 100, 200, 300, 400, 600 and 800 ppm with photosynthetic photon flux density of 2000 μmol m^−2^ s^−1^. RH was approximately 50% in all measurements. A-C_i_ curves at low oxygen conditions were performed by replacement of air by pure N_2_.

### Analyses of Sucrose and Starch Content

Shoots of 15-day-old plant seedlings were harvested. Sucrose and starch were extracted and estimated by using the Sucrose Colorimetric/Spectrophotometric Assay kit (COMIN) and Starch Colorimetric/Spectrophotometric Assay kit (COMIN) according to the manufacturer's instructions, respectively.

### Analyses of Chlorophyll Content and Chlorophyll Fluorescence

Total Chlorophyll content of leaves was extracted with 80 % acetone at 4°C for 24 h in darkness, and then the supernatant was used to measure the absorbance with a spectrophotometer (BeckMan Coulter DU730). Total chlorophyll content was calculated using the following formula:

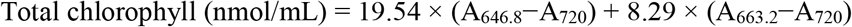

Chlorophyll fluorescence of 15-day-old seedlings was analyzed by using a FluorCam PAM as described in a previous report (Baker, 2008).

### Quantification of Endogenous ABA

0.1 g leaves (fresh weight) of 15-day-old seedlings growing at 200 ppm and 400 ppm were harvested, respectively. Samples were extracted in 750 μl 80:19:1 methanol-H_2_O-acetic acid buffer supplemented with internal standards for 6h with occasional shaking. After centrifugation, the extracts were filtered through a 0.22 μm filter and dried with N_2_ at room temperature. The extracts were dissolved in 200 μl methanol. Quantification of ABA were performed as described (Liu *et al.*, 2012).

### Semi-quantitative PCR and Quantitative Real-time PCR

Total RNA was extracted from seedlings of the wild-type Col-0 and mutants, and cDNA were reverse transcribed. Quantitative RT-PCR was performed by using a Universal SYBR^®^ Green kit and the C1000 Touch Thermal Cycler real-time PCR detection system (Bio-Rad). The *EF1α* (AT5G60390) gene was used as a reference gene for mRNA normalization. Comparative cycle threshold (C_t_) method was used to evaluate the relative gene expression levels. The experiment was repeated three times. The primers used for the expression analysis were listed in Supplemental Table S1.

### GUS Histochemical Analyses

GUS histochemical analyses were carried out on transgenic lines expressing *ProPPC2*∷GUS. Plants at various stages including emerging seedling, 7-day-old seedling, 15-day-old seedling, flowers and siliques were stained in a GUS staining solution and imaged by a Nikon microscope.

### Quantification of Amino Acid and Organic Acid

0.2 g (fresh weight) leaves of 15-day-old seedlings growing at 200 ppm and 400 ppm were harvested, respectively. Tissues were ground and extracted in 8% (w/v) 5-sulfosalicylic acid for 1 h, then were centrifuged. The supernatants were filtered through a 0.22 μm filter. Contents of amino acid were determined by LC-MS/MS methodwith Agilent 1290 Infinity II and Agilent 6460 (Kowalski *et al.*, 2017).

0.2 g (fresh weight) shoots of 15-day-old seedlings growing at 200 ppm and 400 ppm were harvested. Samples were homogenized in 3 ml 7:3 Methanol-Chloroform (−20◻) for 2 h. The water soluble metabolites were extracted from the chloroform phase by adding 2.4 ml H_2_O, after shaking and centrifugation. The upper methanol-H_2_O phase were transferred and dried with N_2_ at room temperature. The extracts were dissolved with 200 μl H_2_O and transferred to 0.45 μm cellulose acetate centrifuge tube filters. The determination of organic acid content was performed by a AB SCIEX QTRAP 6500 Plus LC-MS/MS system as described (Ma *et al.*, 2014).

### Luciferase Assay

Arabidopsis mesophyll protoplasts were isolated from 4 to 6-week-old plants following the method of a previous report (Yoo *et al.*, 2007). 15 μg plasmid DNA was used for PEG-calcium transformation (PEG4000) and the protoplast transformation culture were performed as described (Yoo *et al.*, 2007).

Cellular extracts of Arabidopsis protoplasts after transformation with different constructs for 14 hours were collected for dual-luciferase assays (Hellens *et al.*, 2005). 30 μL of cellular extract was used to detect the firefly and Renilla luciferase activities by a Mithras LB 940 Multimode Microplate Reader. Luciferase activity was normalized to the Renilla activity. All experiments were performed at least three times.

### Accession Numbers

The accession numbers of genes used in this study are available at TAIR (The Arabidopsis Information Resource): *PPC1* (AT1G53310), *PPC2* (AT2G42600), *PPC3* (AT3G14940), *ABI3* (AT3G24650), *ABI4* (AT2G40220), *ABI5* (AT2G36270), *GGAT1* (AT1g23310), *GGAT2* (AT1g70580), *SGAT1* (AT2G13360), *GLDT1* (AT1G11860), *GLDP1* (At4g33010), *SHMT1* (At4g37930).

## Results

### ppc2 mutant seedlings showed growth arrest at low CO_2_ conditions

To investigate whether the *Arabidopsis* PEPCs are involved in plant growth regulation at low CO_2_ conditions, we determined the growth performance of T-DNA insertion lines of plant-type PEPCs, *ppc1* (Salk_088836) (Feria *et al.*, 2016), *ppc2* (Salk_128516) (Shi *et al.*, 2015) and *ppc3* (Salk_143289) (Feria *et al.*, 2016), on sucrose-free 1/2 MS medium at low CO_2_ conditions (200 ppm) for 15 days, as well as the control condition (400 ppm) which was considered as ambient CO_2_ concentration. These single mutants were determined as knockout mutants by our analyses (Fig. 1A and B) and previous studies (Shi *et al.*, 2015; Feria *et al.*, 2016). At ambient CO_2_ conditions, there were no obvious morphological differences among the *ppc* mutants and Col-0 seedlings (Fig. 1C). At low CO_2_ conditions, the cotyledons of Col-0, *ppc1* and *ppc3* turned pale green and no big morphological difference was found among them. Interestingly, *ppc2* mutant plants failed to achieve the same growth state compared to Col-0, *ppc1* and *ppc3* mutant plants (Fig. 1C). *ppc2* mutant plants had small sizes and chlorotic cotyledons, the cotyledons in 87.5% of *ppc2* plants were chlorotic, compared with that of 10% to 20% in Col-0, *ppc1* and *ppc3* plants (Fig. 1D). These results suggest that *PPC2* is required for seedling growth and development at low CO_2_ conditions.

**Fig. 1.**
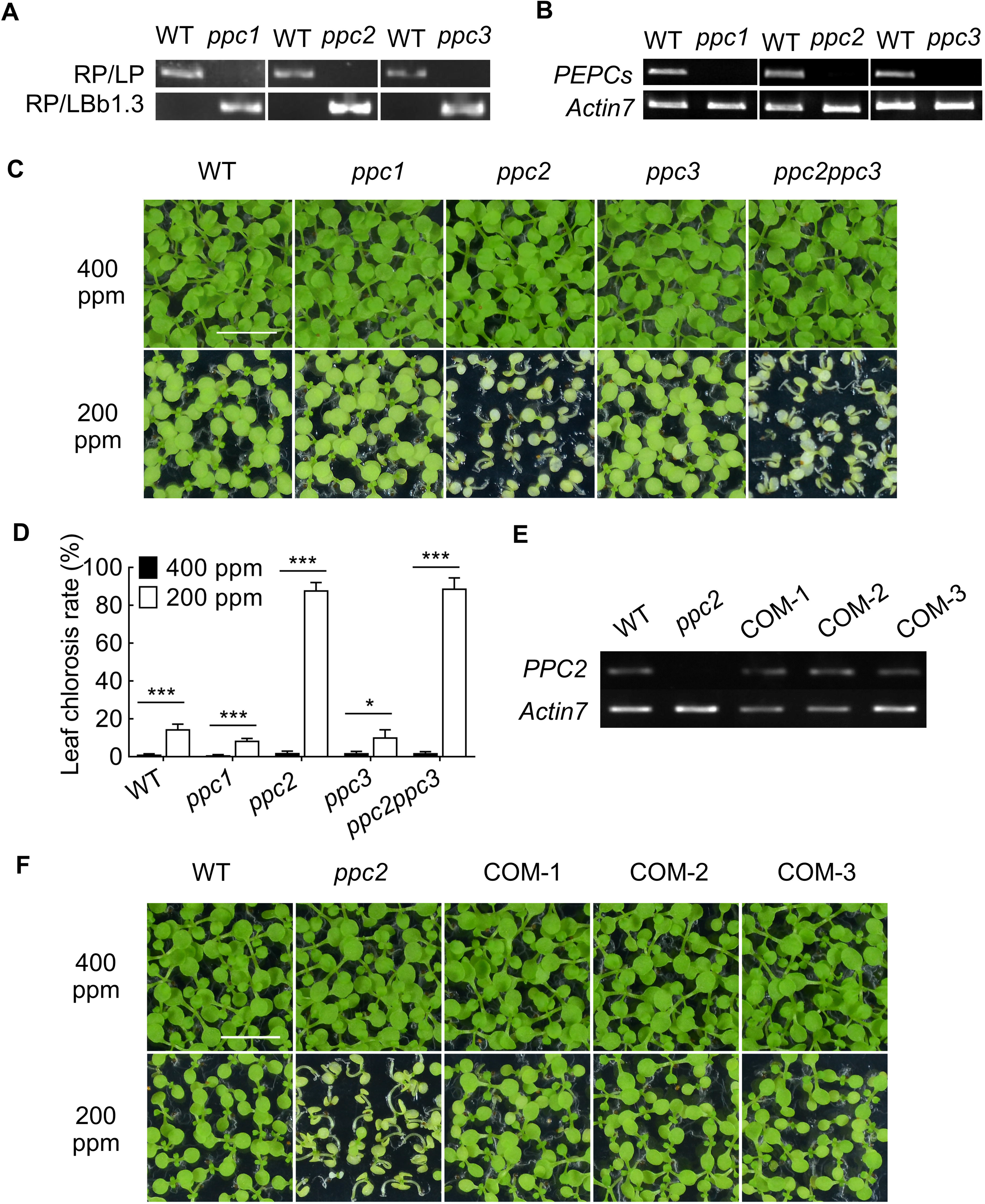
*ppc2* mutant seedlings show growth arrest at low CO_2_ conditions. (A, B) Genotyping analyses (A) and expression levels (B) of *PPC1*, *PPC2* and *PPC3* in their corresponding single mutants of *ppc1*, *ppc2* and *ppc3*. *ACTIN7* (AT5G09810) was used as a control. (C, D) Phenotype (C) and statistic analyses (D) of leaf chlorosis of wild type (WT), *ppc1*, *ppc2*, *ppc3* and *ppc2ppc3* seedlings that growing on sucrose free 1/2 MS medium at 400 ppm or 200 ppm CO_2_ conditions for 15 days. Data shown are mean ± SEM (n = 4). Each replicate had at least 60 seedlings. Asterisks indicate significant differences between genotypes (\**P* < 0.05; \*\*\**P* <0.005 by Student’s t-test; ns, no significant difference). (E) Expression levels of *PPC2* in *PPC2* expressing *ppc2* plants by RT-PCR. RNA was extracted from leaves of 15-day-old seedlings. *ACTIN7* (AT5G09810) was used a control. (F) Growth phenotypes of seedlings of WT, *ppc2* and *PPC2* complementary lines (COM-1, COM-2 and COM-3) growing on sucrose free 1/2 MS medium at 400 ppm or 200 ppm CO_2_ conditions for 15 days. Each replicate had at least 60 seedlings. Bars = 1 cm in C and F.

To determine whether the phenotypes of seedling growth arrest in the *ppc2* mutant are due to the defect of seed germination, the seed-germination rates of these lines were determined at low CO_2_ conditions on sucrose-free 1/2 MS medium. The similar germination rate and status were observed among Col-0, *ppc1*, *ppc2* and *ppc3* (Fig. S1). These results revealed that the growth arrest phenotype of *ppc2* was occurred after germination developmental stage. *AtPPC2* is involved in seedling development at low CO_2_ conditions.

To further reveal whether *PPC1* or *PPC3* has effect on seedling growth at low CO_2_ conditions dependent on *PPC2*, we crossed *ppc1* or *ppc3* with the *ppc2* mutant. We obtained *ppc2ppc3* and *ppc1ppc2* double mutant, but could not get the seeds of *ppc1ppc2* double mutant due to severely growth arrest, consistent with the previous study (Shi *et al.*, 2015). *ppc2ppc3* double mutant showed the similar growth phenotype as *ppc2* single mutant at low CO_2_ conditions and has normal seed germination (Fig. 1C and S1), suggesting *AtPPC3* possibly does not participate in seedling growth regulation at low CO_2_ conditions.

To confirm that *PPC2* was responsible for the phenotype of growth retardation observed in the *ppc2* mutant, the CDS of *PPC2* driven by CaMV 35S promoter was introduced into the *ppc2* mutant. The expression levels of *PPC2* in randomly selected *PPC2*-expressing transgenic *ppc2* plants were recovered to the similar level as in Col-0 by RT-PCR analyses (Fig. 1E). When these lines together with *ppc2* and Col-0 grew at low CO_2_ conditions, the growth arrest and cotyledon chlorotic phenotypes of *ppc2* were all rescued by *PPC2* expression (Fig. 1F). These results demonstrate that the seedling growth arrest is due to the dysfunction of *PPC2* and PPC2 is a major regulator of seedling growth at low CO_2_ conditions.

### PPC2 is low CO_2_ inducible and encodes a major PEPC in Arabidopsis leaves

We next detected the expression levels of these three *PEPCs* at low CO_2_ conditions by real-time PCR analyses, only *PPC2* could be induced by low CO_2_ (Fig. 2A), in agreement with the previous study (Li *et al.*, 2014). To confirm PPC2 is a functional PEPC in Arabidopsis, we measured the total PEPC activity in the *ppc2* mutant and Col-0 at low and ambient CO_2_ conditions. Our results showed that *ppc2* mutant seedlings had lost most of PEPC activity in the leaves at both 400 ppm and 200 ppm CO_2_ conditions (Fig. 2B). Moreover, low CO_2_ treatment increased PEPC activity in Col-0 but not in the *ppc2* mutant leaves (Fig. 2B), further supporting that PPC2 is the major PEPC in *Arabidopsis* leaves and only *PPC2* is in response to low CO_2_.

**Fig. 2.**
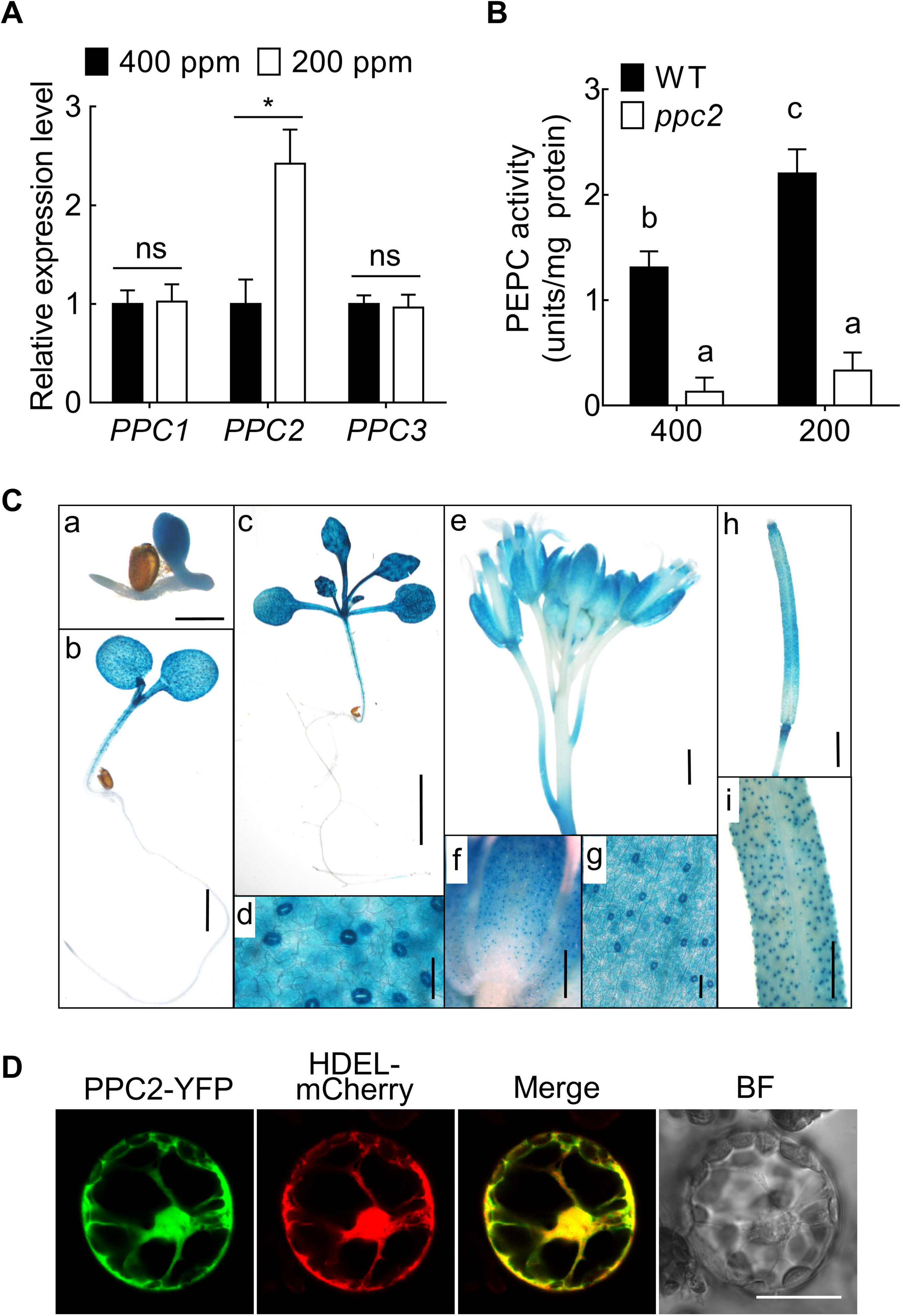
*PPC2* is low CO_2_ inducible and encodes a major PEPC in Arabidopsis leaves. Expression levels of *PPC1*, *PPC2* and *PPC3* in Col-0 15-day-old seedlings that growing on sucrose free 1/2 MS medium at 400 ppm or 200 ppm CO_2_ conditions. Expression levels are expressed relative to that of *EF1α* (AT5G60390). Data shown are mean ± SEM (n = 3). Asterisks indicate significant differences between genotypes (\**P* < 0.05; ns, no significant difference). (B) Total leaf PEPC activity analyzed in 15-day-old wild-type and *ppc2* mutant seedlings at ambient and low CO_2_ conditions. Data shown are mean ± SEM (n = 3). Different letters indicate significant differences using Tukey’s test at *P* ≤ 0.05. (C) Tissue-specific expression of *PPC2* in leaves, flowers and siliques. a to c, GUS activity in leaves of 3-day-old (a), 5-day-old (b) and 15-day-old (c) seedlings. d, guard cells of cotyledon. e and f, GUS activity in calyxes. g, guard cells of calyx. h, GUS activity in siliques. i, guard cells of siliques. Bars = 50 μm in d, g, Bars = 1 mm in a to c, e, f, h and i. (D) Subcellular localization of PPC2-YFP fusion in protoplasts from 4-week-old Col-0. The yellow fluorescence from PPC2-YFP was shown in green and merged with red fluorescence from the HDEL-mCherry fusion using confocal microscopy. BF, bright field. YFP, yellow fluorescent protein. Bar = 20 μm.

We then determined its spatial expression patterns by expressing *GUS* reporter gene driven by *PPC2* promoter in Col-0. GUS staining showed that *PPC2* was mainly expressed in leaves, hypocotyl, flowers and siliques, but not or very weak in roots (Fig. 2C). We also found *PPC2* was highly expressed in guard cells. We were also interested to know the subcellular localization of PPC2, *35S∷PPC2-YFP* construct was transformed into the protoplasts of Col-0. YFP fluorescence of PPC2-YFP fusion protein revealed its localization in the cytoplasm, nucleus and also endoplasmic reticulum (Fig. 2D). Taken together, these results demonstrate a specific role of *PPC2* in regulation of plant growth at low CO_2_.

### ppc2 seedlings showed reduced carbon assimilation at low CO_2_ conditions

To confirm whether the growth arrest and cotyledon chlorosis of *ppc2* mutant at low CO_2_ is due to the defect of carbohydrate accumulation, we detected starch content by iodine staining and quantification in 15-day-old *ppc2* and Col-0 seedlings at the end of the illumination period (22:00 pm) and darkness period (06:00 am) at different CO_2_ conditions. At ambient CO_2_ conditions there was no obvious difference in starch accumulation between *ppc2* and Col-0 plants (Fig. S2A and B), however, at low CO_2_ conditions the starch accumulation was significantly reduced in the *ppc2* cotyledons at both time points (Fig. S2A and B). Because in plant cells, starch is synthesized at daytime and degraded at night (darkness). We then measured sucrose content of *ppc2* and Col-0 at these conditions. The sucrose content was lower in the *ppc2* seedlings than in Col-0 at either low or ambient CO_2_ conditions (Fig. S2B). Moreover, the synthetic substrates of starch and sucrose, such as G6P, F6P, G1P, ADPG, UDPG and Suc6P were decreased in *ppc2* at low CO_2_ conditions, but were similar at ambient CO_2_ conditions compared to Col-0 (Fig. S2B), consistent with the starch and sucrose content in *ppc2*. These findings suggest that *ppc2* mutation led to the reduced photosynthetic carbohydrate accumulation at low CO_2_ conditions.

To test the reduction of photosynthetic carbohydrates in *ppc2* is caused by the reduced CO_2_ assimilation, we measured chlorophyll and carotenoid contents in these seedlings. At ambient CO_2_ conditions, there were no significant differences in total chlorophyll and carotenoid contents (Fig. 3A and B). At low CO_2_ conditions both chlorophyll and carotenoid were greatly reduced in *ppc2*. We next detected the maximal quantum yield of PSII (*F*_*v*_/*F*_*m*_) in *ppc2* and Col-0 seedlings growing at 200 ppm and 400 ppm CO_2_ conditions by chlorophyll florescence detector FluorCam FC800. At ambient CO_2_ conditions there was no difference in *F*_*v*_/*F*_*m*_ between Col-0 and *ppc2*. Low CO_2_ treatment greatly reduced *F*_*v*_/*F*_*m*_ in both Col-0 and *ppc2*, however a remarkable decrease was observed in *ppc2* than in Col-0 (Fig. 3C and D). These results suggest *PPC2* is involved in photosynthesis regulation at low CO_2_.

**Fig. 3.**
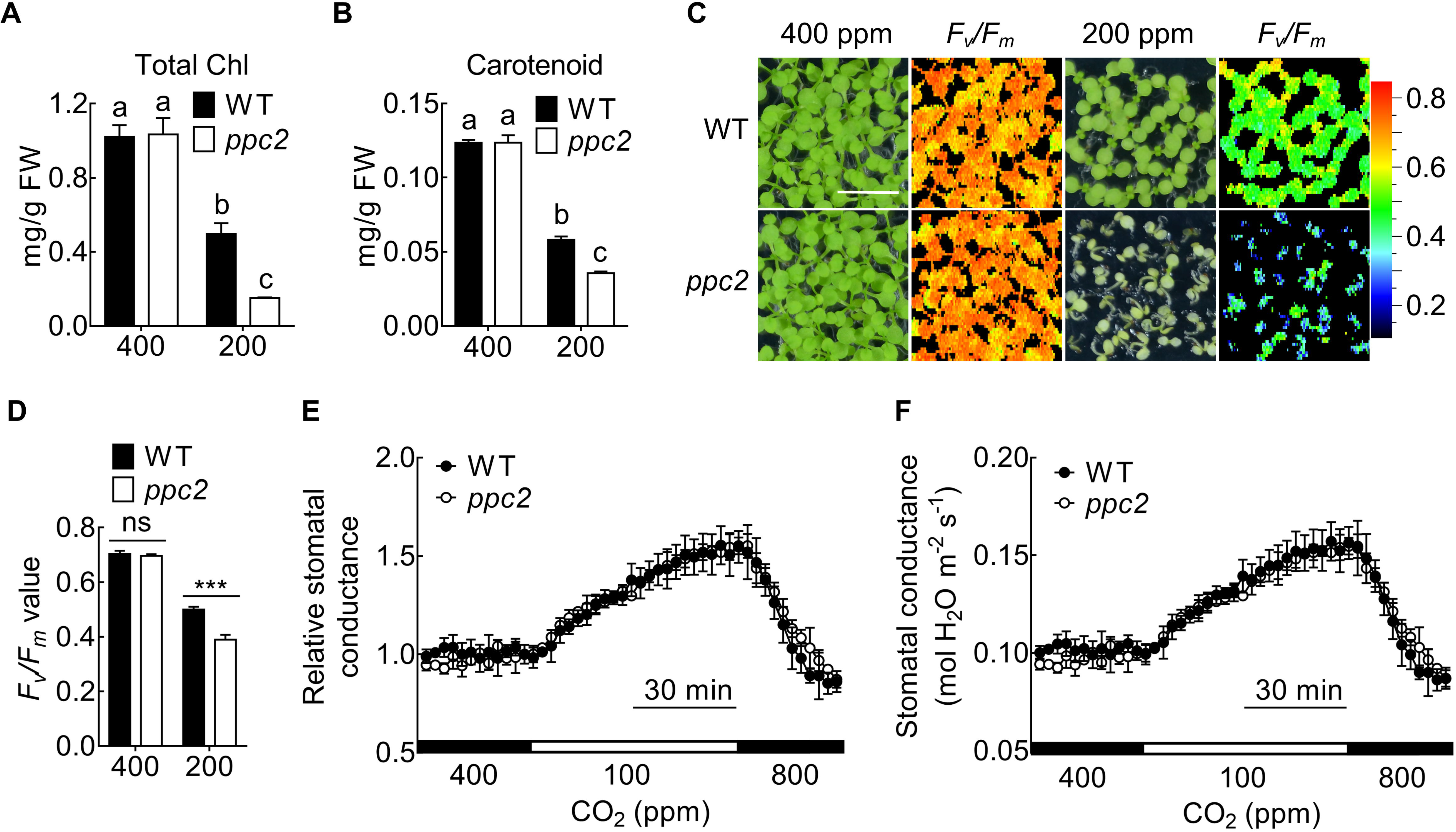
Carbon assimilation is reduced in the *ppc2* mutant at low CO_2_ conditions. (A, B) Chlorophyll content (A) and carotenoid content (B) in 15-day-old seedlings of wild type (WT) and *ppc2* mutant at 400 ppm and 200 ppm CO_2_ conditions. Data shown are mean ± SEM (n = 3). (C) *F*_*v*_/*F*_*m*_ was monitored by Closed FluorCam FC800 in WT and *ppc2* mutant seedlings growing at 400 ppm or 200 ppm CO_2_ conditions for 15 days. (D) *F*_*v*_/*F*_*m*_ value comparison between WT and *ppc2* mutant. Data shown are mean ± SEM (n = 3). (E, F) Time-resolved stomatal conductance in response to CO_2_ changes in WT and *ppc2* mutant plants. (E) Data shown in (F) were normalized. Different letters indicate significant differences using Tukey’s test at *P* ≤ 0.05. Asterisks indicate significant differences between genotypes (\*\*\**P* <0.005 by Student’s t-test; ns, no significant difference).

Because *PPC2* is highly expressed in guard cell (Fig. 2C), it is important to confirm if the reduced carbon assimilation rate at low CO_2_ conditions was induced by the compromised low CO_2_-induced stomatal opening. The CO_2_ shifts from 400 to 100 ppm triggered dramatic increase in stomatal conductance in both Col-0 and *ppc2* mutant plants. However, there were no big difference in stomatal response to low CO_2_ between the *ppc2* mutants and Col-0 (Fig. 3E and F), indicating minor role of *PPC2* in regulating stomatal opening, and also suggesting the reduced carbon assimilation in *ppc2* at low CO_2_ conditions is not caused by reduced CO_2_ uptake but by CO_2_ utilization.

### Addition of exogenous sucrose or malate greatly rescued the seedling growth arrest of ppc2 at low CO_2_ conditions

Because of the reduced photosynthetic carbohydrate accumulation and carbon assimilation in the *ppc2* mutant at low CO_2_ conditions, we would like to know whether application of sucrose, which is a photosynthetic product, could rescue the reduced seedling growth of *ppc2* mutant at low CO_2_ conditions. Exogenous sucrose (25 mM) treatment completely recovered the seedling growth retardation and cotyledon chlorosis in *ppc2* at low CO_2_ conditions (Fig. 4A).

**Fig. 4.**
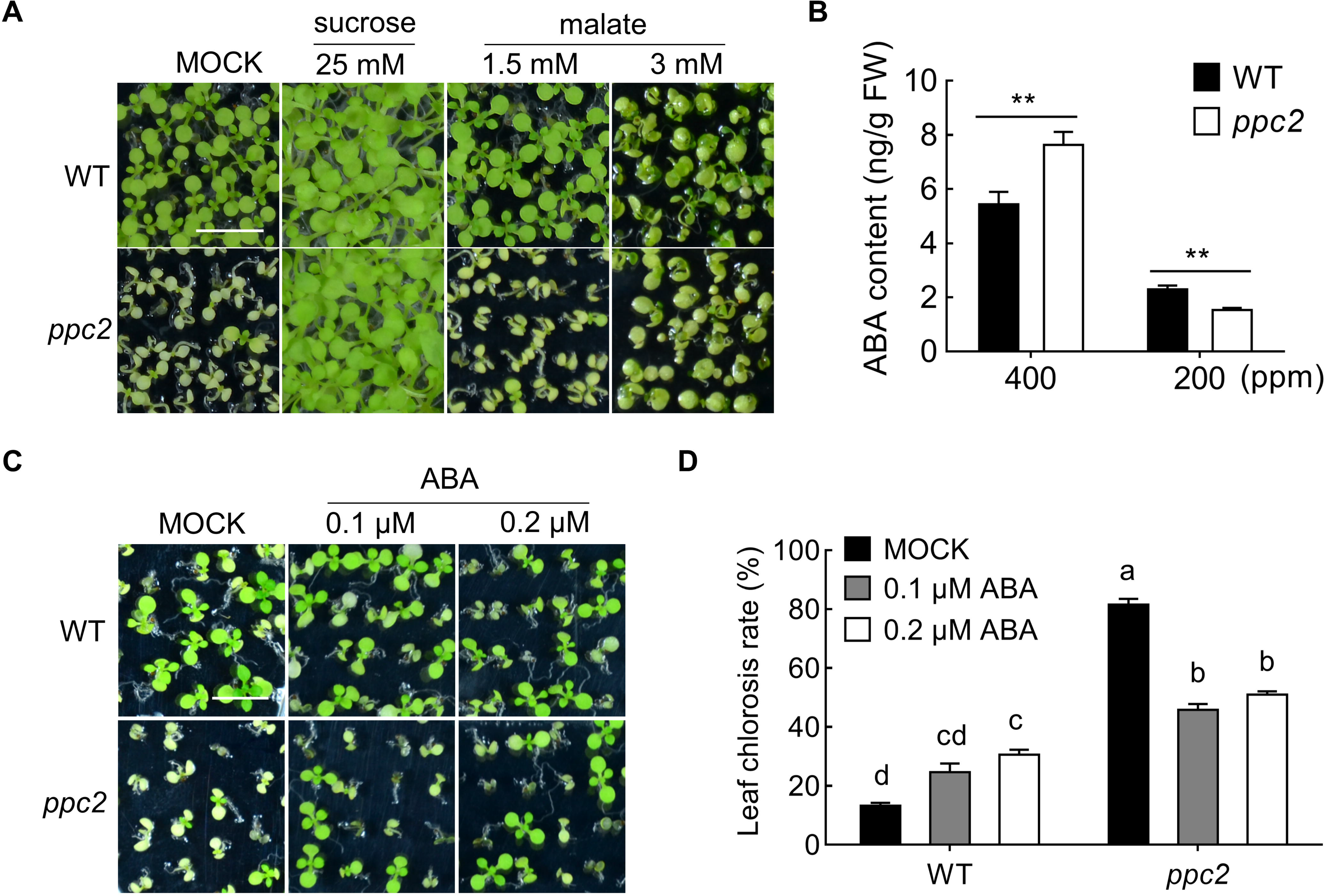
Exogenous ABA greatly rescues the seedling growth arrest of *ppc2* at low CO_2_ conditions. (A) Phenotype of 15-day-old seedlings of wild type (WT) and *ppc2* mutant at 200 ppm CO_2_ on sucrose free 1/2 MS medium, supplemented with mock (ddH_2_O), 25 mM sucrose, 1.5 mM malate or 3 mM malate. Each replicate had at least 60 seedlings. (B) ABA content in WT and *ppc2* mutant seedlings growing at different CO_2_ conditions. Data shown are mean ± SEM (n = 3). FW, fresh weight. (C, D) Phenotypes (C) and statistic analyses (D) of leaf chlorosis rate of 15-day-old WT and *ppc2* mutant seedlings growing at 200 ppm CO_2_ on sucrose free 1/2 MS medium with mock (ddH_2_O), 0.1 μM ABA, 0.2 μM ABA, respectively. Data shown are mean ± SEM (n = 4). Each replicate had at least 60 seedlings. Asterisks indicate significant differences between genotypes (\*\**P* <0.01 by Student’s t-test). Different letters indicate significant difference using Tukey’s test at *P* ≤ 0.05. Bar = 1 cm in A and C.

PEPC catalyzes the synthesis of oxaloacetate (OAA) from PEP and HCO_3_^−^. The product OAA is rapidly converted into malate by malate dehydrogenase. Our primary metabolic analyses showed malate was reduced when *PPC2* was mutated (Fig. S2B), and their differences between *ppc2* mutant and Col-0 were increased at low CO_2_ conditions. We then explored whether the growth-arrest phenotype of *ppc2* mutant at low CO_2_ was caused by malate defect. Application of 1.5 mM malate didn't have the effect on growth of both genotypes and could not eliminate the cotyledon phenotype of *ppc2* (Fig. 4A), however the growth-arrest phenotype of *ppc2* was greatly relieved by addition of 3 mM malate compared to control group at our growth conditions. These results support the function of *AtPPC2* in both carbon assimilation and metabolic pathway.

### Exogenous ABA partially rescued the seedling growth arrest of ppc2 at low CO_2_ conditions

ABA is synthesized from C_40_ oxygenated carotenoids (Ruiz-Sola and Rodriguez-Concepcion, 2012) and in the *ppc2* mutant carotenoid content was seriously reduced (Fig. 3B). To determine whether ABA biosynthesis is blocked in *ppc2* at low CO_2_ conditions. We quantified ABA level in 15-day-old *ppc2* and Col-0 seedlings that growing at low and ambient CO_2_ conditions by UFLC-ESI-MS (Liu *et al.*, 2012). Interestingly, ABA content in the *ppc2* mutant was greatly reduced at low CO_2_ conditions but increased at atmospheric CO_2_ conditions compared to Col-0 (Fig. 4B). These results demonstrate that the reduced ABA level may contribute to the seedling growth arrest in *ppc2* at limiting CO_2_ conditions.

To prove this, we added low concentrations of exogenous ABA (0.1 or 0.2 μM) to sucrose-free 1/2 MS medium plate to see the growth performance of Col-0 and *ppc2*. In Col-0, ABA treatment inhibited the growth and increased the ratio of chlorotic seedlings, while in the *ppc2* mutant ABA treatment largely recovered the cotyledon chlorotic and growth arrest phenotypes (Fig. 4C and D). Taken together, our results demonstrate that *ppc2* mutation affects ABA biosynthesis and the seedling growth-arrest phenotype of *ppc2* mutant is at least partly due to the reduced ABA biosynthesis.

### abi5 showed reduced F_v_/F_m_ and expression of ABI5 greatly rescued the growth arrest of ppc2 seedlings at low CO_2_ conditions

It has been reported that *ABI3*, *ABI4* and *ABI5*, downstream targets of ABA signaling pathway, are required for the ABA modulation of seed germination and postgermination development (Finkelstein and Lynch, 2000; Lopez-Molina *et al.*, 2001). To determine whether ABI transcription factors function in the seedling growth arrest in the *ppc2* mutant at low CO_2_ conditions, we firstly checked their expression levels by qPCR. Low CO_2_ treatment induced the expression levels of all these three *ABIs* in Col-0. The expression of *ABI5* at low CO_2_ conditions was greatly suppressed in the *ppc2* mutant (Fig. 5A), however the expression levels of *ABI3* and *ABI4* were comparable in Col-0 and *ppc2* seedlings at both ambient and low CO_2_ conditions. Furthermore, exogenous ABA could rescue the *ABI5* expression at low CO_2_ conditions in the *ppc2* mutant (Fig. 5B). These results indicate the reduced expression level of *ABI5* might be a major cause of reduced seedling growth in *ppc2* at low CO_2_ conditions. To confirm this and reveal the role of *ABI5* in seedling growth at low CO_2_ conditions, we determined the growth performance of *abi5-1* mutant and wild type Ws plants at low CO_2_ conditions on sucrose-free medium. No obvious morphological differences were observed between Ws and *abi5-1* seedlings (Fig. 5C). However, *F*_*v*_/*F*_*m*_ level was significantly reduced in the *abi5-1* mutant compared to that in Ws at low CO_2_ conditions (Fig. 5D), indicating mutation of *ABI5* led to reduced maximum quantum efficiency at low CO_2_ conditions. We further overexpressed *ABI5* in the *ppc2* mutant plant driven by the constitutive cauliflower 35S promoter. Expression of *ABI5* greatly rescued the early seedling growth arrest phenotype of *ppc2* in three randomly selected transgenic lines at low CO_2_ conditions (Fig. 5E and F, Fig. S3).

**Fig. 5.**
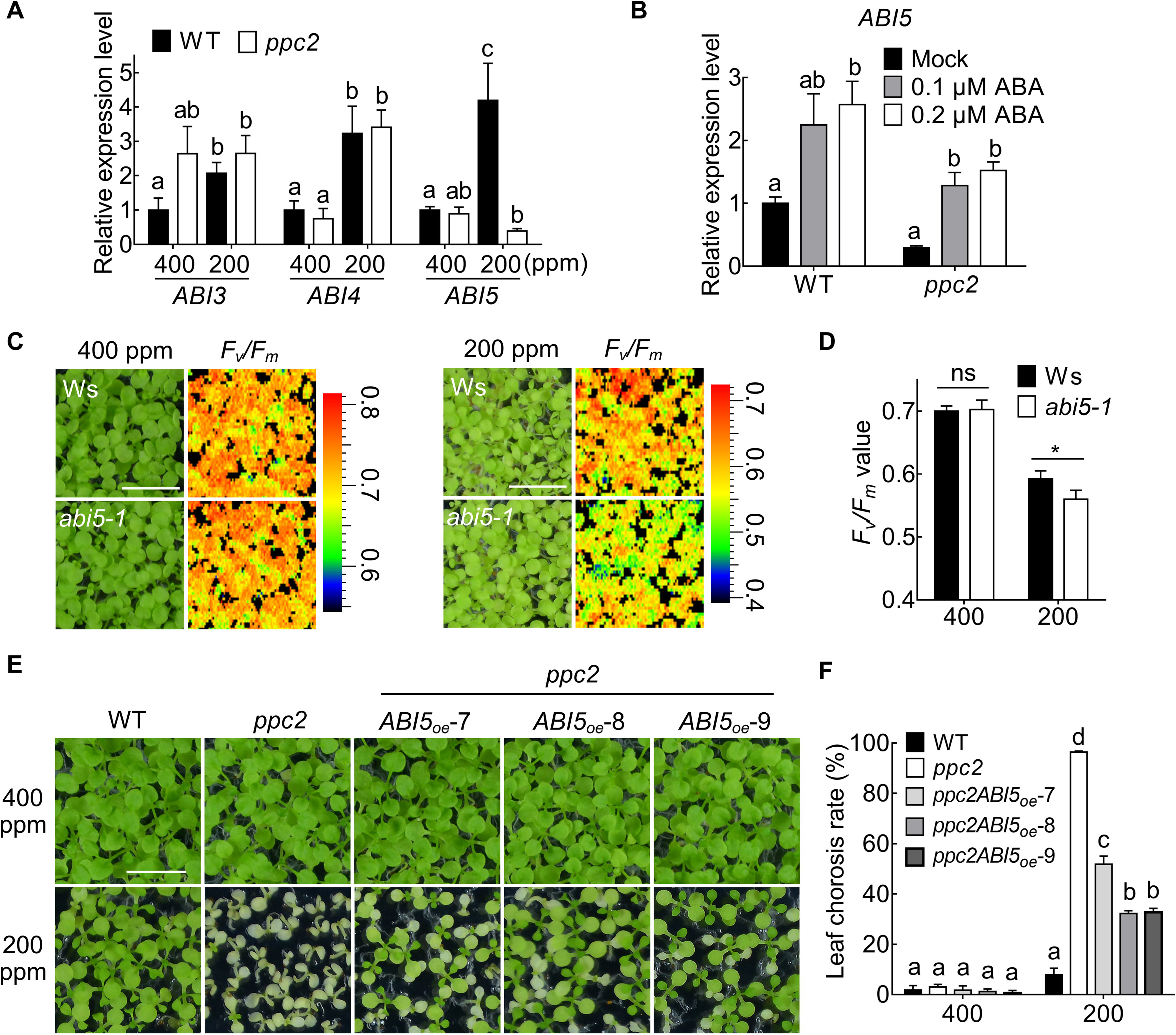
*abi5* has reduced *F*_*v*_/*F*_*m*_ and expression of *ABI5* rescued the growth arrest of *ppc2* at low CO_2_ concentration. (A) Relative expression levels of *ABI3*, *ABI4* and *ABI5* to *EF1α* (AT5G60390) in wild type (WT) and *ppc2* mutant growing on sucrose free 1/2 MS medium at 400 ppm or 200 ppm CO_2_ conditions for 15 days. Data are shown as mean ± SEM (n= 3). (B) *ABI5* expression levels in *ppc2* and WT seedlings that treated with mock (ddH_2_O), 0.1 μM ABA, 0.2 μM ABA at 200 ppm conditions, respectively. Data shown are mean ± SEM (n = 3). Different letters indicate significant differences using Tukey’s test at *P* ≤ 0.05. (C) *F*_*v*_/*F*_*m*_ monitored by Closed FluorCam FC800 in Ws and *abi5-1* mutant seedlings growing at 400 ppm or 200 ppm CO_2_ conditions for 15 days. (D) Maximum photosynthetic yields (*F*_*v*_/*F*_*m*_) of Ws and *abi5-1* mutant at different CO_2_ conditions. Data shown are mean ± SEM (n = 3). Asterisks indicate significant differences between genotypes (\**P* < 0.05 by Student’s t-test; ns, no significant difference). (E, F) Phenotype (E) and statistic analyses (F) of leaf chlorosis rate of WT, *ppc2* mutant and *ABI5* expressing *ppc2* transgenic lines (*ppc2ABI5*_oe_-7, *ppc2ABI5*_oe_-8, *ppc2ABI5*_oe_-9). Data shown are mean ± SEM (n = 3). Each replicate had at least 60 seedlings. Different letters indicate significant difference using Tukey’s test at P ≤ 0.05. Bar = 1cm in C and E.

### Photorespiratory intermediates were increased in ppc2 at low CO_2_ conditions

It is suggested that PEPC activity is linked to photorespiration by supplying malate into the TCA cycle to sustain glutamate and glutamine metabolism (Masumoto *et al.*, 2010; Shi *et al.*, 2015). We then determined amino acids levels in *ppc2* and Col-0 at both ambient and low CO_2_ conditions. Our analyses showed that *ppc2* mutant had reduced glutamic acid and increased glutamine, which led to increased ratio of Gln to Glu (Fig. 6A and B), increased level of β-Alanine and Arginine, and decreased level of alanine, asparatic acid and proline (Fig. S4). Glycine and serine levels have been recognized as indicators of carbon flux through photorespiration and higher ratios of glycine/serine indicate high photorespiration (Novitskaya *et al.*, 2002). Low CO_2_ concentrations would increase photorespiration. *ppc2* mutant had higher contents of photorespiratory intermediates glycine and serine at photorespiratory (low CO_2_) conditions (Fig. 6A) and higher glycine content at 400 ppm CO_2_ concentration (Fig. S4), which were consistent with the reduced photosynthesis and growth arrest at low CO_2_ conditions (Figs. 3E and 4A). We found low CO_2_ (photorespiratory conditions) triggered an increase in glycine/serine ratio both in the *ppc2* mutant and Col-0 in contrast to ambient CO_2_, however no significant differences in glycine/serine ratio were found between them (Fig. 6C). At ambient CO_2_ conditions *PPC2* didn’t have significant effect on amino acid and organic acid content, only glycine, valine and tyrosine were slightly increased in *ppc2* (Fig. S2B and S4). These results suggest *PPC2* functions in both primary metabolism and photorespiratory metabolism at photorespiratory low CO_2_ conditions through modulation of carbon/nitrogen balance.

**Fig. 6.**
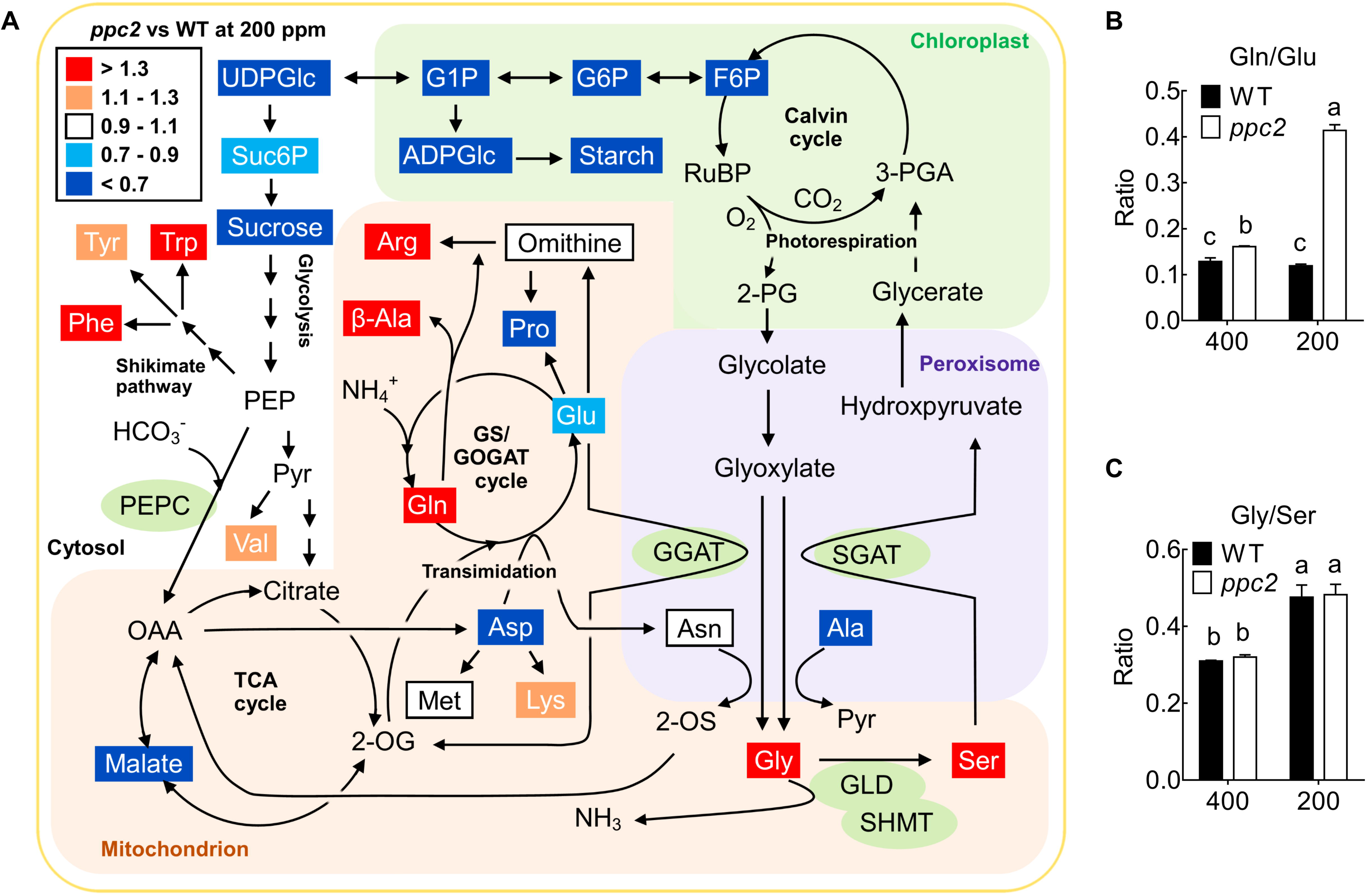
*ppc2* mutant has reduced carbon assimilation rate at different CO_2_ concentrations. (A) Level changes in leaf metabolites between *ppc2* mutant and wild type (WT) seedlings at 200 ppm CO_2_ conditions. PEPC, phosphoenolpyruvate carboxylase. GGAT, glutamate:glyoxylate aminotransferase. GLD, glycine decarboxylase, including GLDP, GLDT and GLDH. SHMT, serine hydroxymethyltransferase. SGAT, serine:glyoxylate aminotransferase. RuBP, ribulose-1,5-disphosphate. F6P, fructose 6-phosphate. G6P, glucose 6-phosphate. G1P, glucose 1-phosphate. UDPGlc, UDP-glucose. ADPGlc, ADP-glucose. Suc6P, sucrose 6-phosphate. Pyr, pyruvate. 2-OS, 2-oxosuccinamate. OAA, oxaloacetate. PEP, phosphoenlpyruvate. (B) The ratio of glutamine to glutamic acid in 15-day-old WT and *ppc2* seedlings at different CO_2_ conditions. Data shown are mean ± SEM (n = 3). (C) The ratio of glycine to serine in 15-day-old WT and *ppc2* seedlings at different CO_2_ conditions. Data shown are mean ± SEM (n = 3). Different letters indicate significant difference using Tukey’s test at P ≤ 0.05.

### ABI5 regulates the expression levels of photorespiratory enzymes

To further explore whether the higher levels of serine and glycine in *ppc2* at low CO_2_ conditions are caused by the reduced levels of those enzymes that synthesize glycine and serine during photorespiration, we checked the expression levels of *GGAT1*, *GGAT2*, *SGAT1*, *GLDP1*, *GLDT1* and *SHMT1* (Peterhansel *et al.*, 2010) in *ppc2* at both low and ambient CO_2_ conditions. In the photorespiratory pathway GGAT (glutamate:glyoxylate aminotransferase) transfers -NH_3_^+^ from glutamate into glyoxylate to generate glycine and SGAT1 (serine:glyoxylate aminotransferase, TAIR abbreviation: AGT) transfers -NH_3_^+^ from Ser, Ala and Asn into glyoxylate to generate glycine. GLDP1 and GLDT1 were enzyme components of glycine decarboxylase complex, catalyzing glycine into CH_2_-THF. The expression levels of these genes except *GGAT2* were significantly reduced in the *ppc2* mutant at low CO_2_ conditions (Fig. 7A). Among them, *GGAT1* and *SGAT1* were slightly induced by low CO_2_ treatment in Col-0 (Fig. 7A). The transcript of *SHMT1* in the *ppc2* mutant was significantly reduced at both ambient and low CO_2_ conditions (Fig. 7A).

**Fig. 7.**
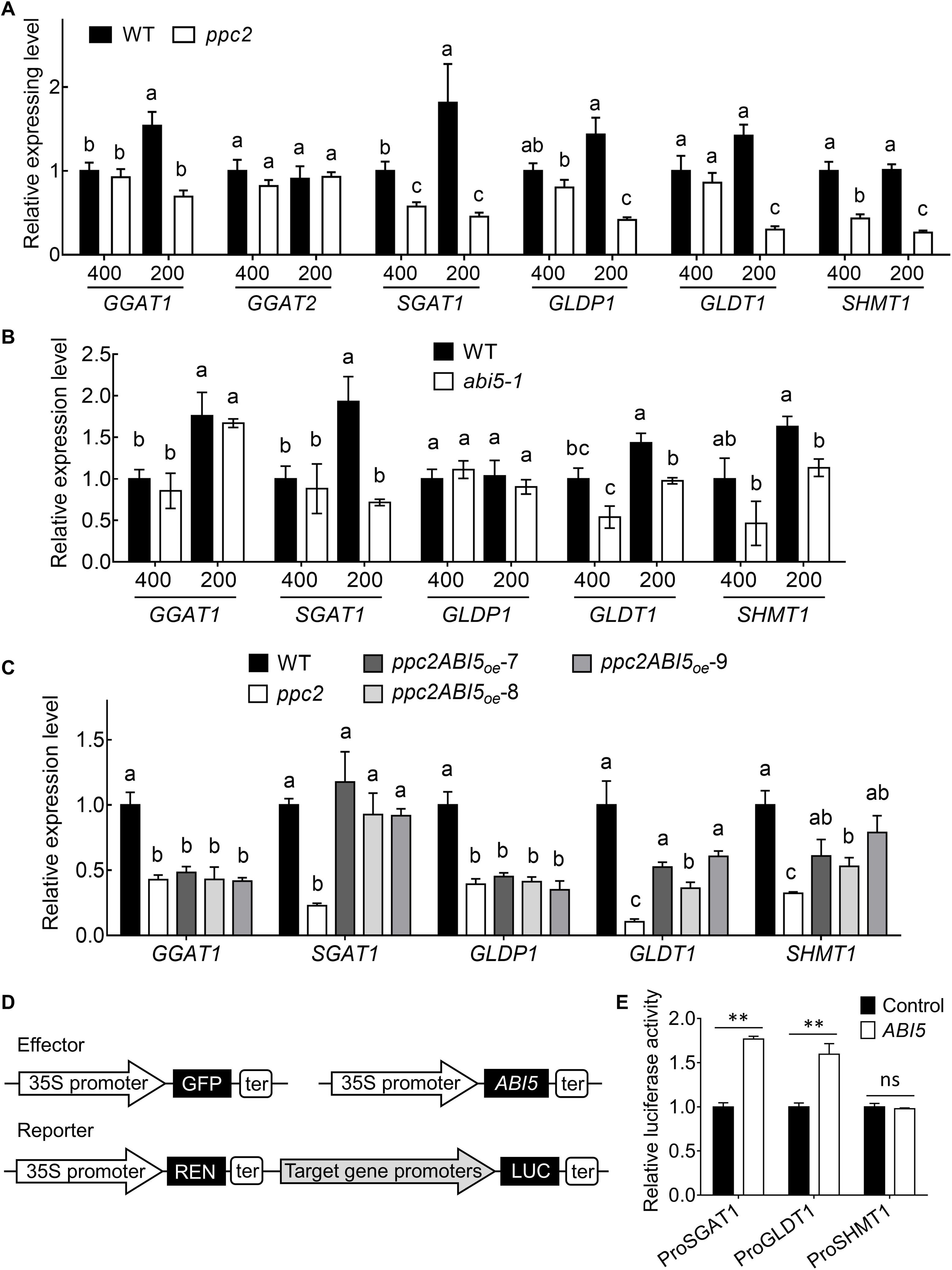
*ABI5* regulates the expression levels of major photorespiratory enzymes related to glycine and serine synthesis and metabolism. (A) Expression levels of photorespiratory enzyme genes in wild type (WT) and *ppc2* mutant leaves. RNAs were extracted from the leaves of 15-day-old seedlings. *EF1α* (AT5G60390) was used as an internal control. Data shown are mean ± SEM (n = 3). (B) Expression levels of photorespiratory genes in WT, *ppc2* mutant and *ABI5* expressing *ppc2* plants (*ppc2ABI5*_oe_-7, *ppc2ABI5*_oe_-8, *ppc2ABI5*_oe_-9). RNAs were extracted from leaves of the 15-day-old seedlings at different CO_2_ condition. *EF1α* (AT5G60390) was used an internal control. Data shown are mean ± SEM (n = 3). Different letters indicate significant differences using Tukey’s test at *P* ≤ 0.05. (B) Expression levels of photorespiratory enzyme genes in WT (Ws) and *abi5-1* mutant leaves. RNAs were extracted from the leaves of 15-day-old seedlings. *EF1α* (AT5G60390) was used as an internal control. Data shown are mean ± SEM (n = 3). (D) A schematic representation of dual luciferase reporter system. *ABI5* or GFP (control) driven by CaMV 35S as effector was co-transformed with reporter that containing 35S driving REN (Renilla Luciferase) and the tested promoter of photorespiratory enzyme genes driving LUC (Firefly Luciferase) expression. (E) Dual-luciferase reporter assays showed the transcriptional activation of *ABI5* on the promoters of *SGAT1*, *GLDT1* and *SHMT1*. LUC values were normalized to REN. Data shown are mean ± SEM (n = 3). Asterisks indicate significant differences between genotypes (\*\**P* < 0.01 by Student’s t-test; ns, no significant difference).

ABI5 is a transcription factor that directly binds to the promoter regions of its targets to activate their expressions. *ABI5* expression was reduced in *ppc2* and *ABI5* overexpression in *ppc2* recovered the reduced growth arrest at photorespiratory low CO_2_ conditions, we thus hypothesized that ABI5 might regulate the expression levels of these enzymes that function in glycine and serine production, such as *GGAT1*, *SGAT1*, *GLDP1*, *GLDT1* and *SHMT1*. We interestingly found there were several ABREs, ABI5 binding cis-element, in the promoter regions of these five genes (Fig. S5). The expressions of *SGAT1*, *GLDT1* and *SHMT1* were reduced in *abi5-1* mutant at low CO_2_ conditions (Fig. 7B), suggesting these three genes could be the targets of ABI5. We interestingly found the expression levels of *SGAT1*, *GLDT1* and *SHMT1* in three randomly selected independent transgenic lines were completely or greatly recovered (Fig. 7C). We then performed dual luciferase assays to determine the ABI5 activation of the promoters of *SGAT1*, *GLDT1* and *SHMT1* that drive LUC expression in Arabidopsis mesophyll cell protoplasts. ABI5 expression greatly activated the promoters of *SGAT1* and *GLDT1*, but could not activate the promoter of *SHMT1* (Fig. 7D and E). These results demonstrate that *SGAT1* and *GLDT1* could be the direct targets and *SHMT1* was an indirect target of ABI5, and ABI5 regulates photorespiration by modulating the expression levels of photorespiratory enzymes.

### A-C_i_ curve at photorespiratory low CO_2_ conditions was reduced in the ppc2 mature plants

Recently, a potential role for PEPC in C_3_ plant metabolism at high photorespiratory (low CO_2_/high O_2_) conditions has been proposed (Abadie and Tcherkez, 2019; Tcherkez and Limami, 2019). Here we also found that PPC2 is involved in seedling development by modulation of photorespiratory metabolism at low CO_2_ conditions. To further investigate the function of *Arabidopsis PPC2* in photorespiratory conditions, we determined the CO_2_ assimilation rate under different C_i_ conditions in adult leaves of *ppc2* and Col-0 under air condition. The *ppc2* mutant exhibited a reduced CO_2_ assimilation rate at low CO_2_ (50-400 ppm) concentrations in the initial part of the A-C_i_ curve compared with Col-0, while exhibiting similar rate with Col-0 at higher CO_2_ (400-800 ppm) concentrations (Fig. 8A). In addition, the maximum photosynthetic electron transport rates of Col-0 and *ppc2* mutant plants which were computed from A-C_i_ curves exhibited no significant difference (Fig. 8B), indicating the photosynthetic capacity of the *ppc2* mutant was not changed. The reduce in the initial A-C_i_ curve was recovered when the measurements were performed under very low oxygen conditions that restricted photorespiration (Fig. 8C). These results indicate the altered CO_2_ assimilation of *ppc2* mutant at low CO_2_ conditions was associated with the simultaneous high photorespiratory lost in plant. Moreover, *PPC2* expression rescued the reduced photosynthetic rate in response to low C_i_ changes in *ppc2* (Fig. 8D). Together with that PEPC is an important enzyme in the glycolytic pathway and links with photorespiration and respiration in plant, our results suggest that *PPC2* is involved in photorespiration at relatively low CO_2_ conditions. The phenotype of *ppc2* is, at least partially, due to the reduced net carbon assimilation, which may be resulted from the low capacity to utilize the photorespiratory metabolites at relatively low CO_2_ conditions when *PPC2* is mutated.

**Fig. 8.**
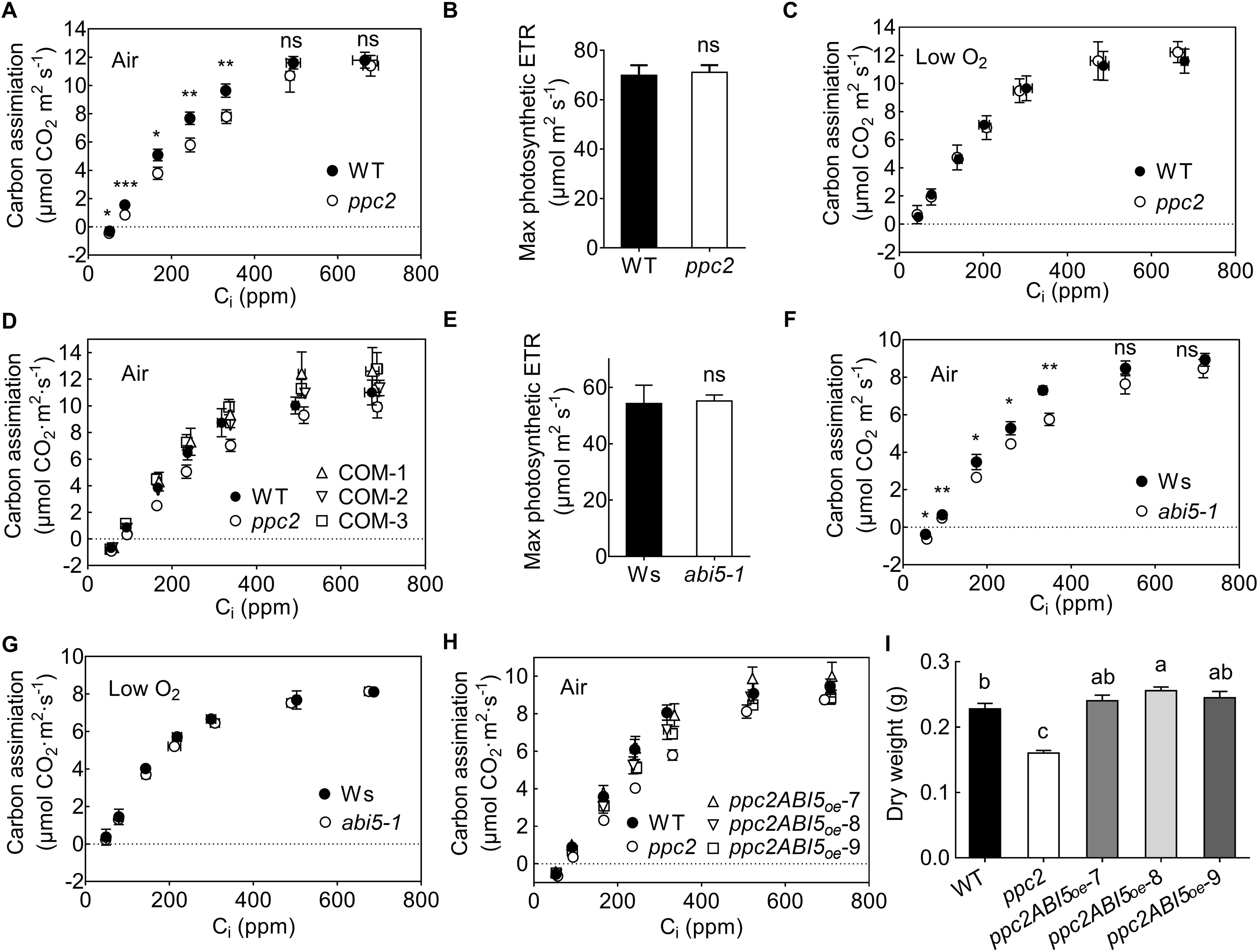
*abi5* mutant shows reduced carbon assimilation and expression of *ABI5* rescues the carbon assimilation of *ppc2* mutant at photorespiratory low CO_2_. (A, C) A-C_i_ curves of 30-day-old wild type (WT) and *ppc2* mutant plants under ambient air conditions (A) or low oxygen condition (C). The light intensity for measurement was set at 2000 μmol m^−2^ s^−1^. Data shown are mean ± SEM (n = 3). (B) Maximum photosynthetic electron transport rate (ETR) of 30-day-old wild type and *ppc2* mature plants. Data shown are mean ± SEM (n = 5). (D) Expression of *PPC2* complements the reduced A-C_i_ curves on rosette leaves of *ppc2* mutant growing at 400 ppm conditions (COM-1, COM-2 and COM-3 are *PPC2* complementary lines). Data shown are mean ± SEM (n = 3). (E) Maximum photosynthetic electron transport rate (ETR) of 30-day-old Ws and *abi5-1* mutant plants. Data shown are mean ± SEM (n = 5). (F, G) A-C_i_ curves of *abi5-1* and Ws rosette leaves under air (F) or low oxygen (G) conditions for 30 days. Data shown are mean ± SEM (n = 3). (H) *ABI5* expression complements the reduced A-C_i_ curves of *ppc2* plants that growing at 400 ppm conditions. Data shown are mean ± SEM (n = 3). (I) Dry weight of WT, *ppc2* and *ABI5* overexpression *ppc2* transgenic lines (*ppc2ABI5*_oe_-7, *ppc2ABI5*_oe_-8, *ppc2ABI5*_oe_-9) growing at ambient CO_2_ conditions. Data shown are mean ± SEM (n = 6). Asterisks indicate significant differences between genotypes (\**P* < 0.05; \*\**P* < 0.01; \*\*\**P* <0.005 by Student’s t-test; ns, no significant difference). Different letters indicate significant differences using Tukey’s test at *P* ≤ 0.05.

### Overexpression of ABI5 rescued the reduced CO_2_ assimilation at photorespiratory low CO_2_ conditions in the ppc2 mature plants

We also determined the A-C_i_ curves of *abi5-1* 30-day-old plants at air and low oxygen conditions, respectively. When measurements were performed under air condition, mutation of *ABI5* greatly reduced the CO_2_ assimilation rate at low CO_2_ concentrations (50-400 ppm), but which was similar at high CO_2_ concentrations (600-800 ppm) compared to Col-0 (Fig. 8F). The max photosynthetic ETR of Ws and *abi5-1* inferred from A-C_i_ curves in air condition were similar (Fig. 8E). In low oxygen conditions, the slopes of A-C_i_ curves of *abi5-1* and Ws had no significant difference between *abi5* and Ws (Fig. 8G). These results demonstrate that *ABI5* is involved in photorespiratory metabolism pathway.

We next determined the A-C_i_ curves of *ABI5*-expressing *ppc2* plants growing at ambient CO_2_ conditions. The reduced CO_2_ assimilation of *ppc2* at low CO_2_ conditions was totally rescued by *ABI5* expression (Fig. 8H). Moreover, *ABI5*-expressing not only restored the reduced dry-weight biomass of *ppc2* mutant but also had some increase at ambient conditions (Fig. 8I).

## Discussion

### PPC2 is essential for plant growth at low CO_2_ conditions

CO_2_ is the major source for photosynthesis and is pivotal for plant growth. High CO_2_ often increases plant growth and reproduction, whereas low CO_2_ decrease plant growth by changing physiological responses (e.g. water use efficiency), reducing biomass production and delaying development (Gerhart and Ward, 2010). However, the mechanisms are still unclear. Here we report that Arabidopsis *PPC2*, encoding a PEPC that is involved in plant primary metabolism for producing C4-dicarboxylic acids, is essential plant growth at low CO_2_ conditions. *ppc2* seedlings showed chlorosis and arrested growth at low CO_2_ (200 ppm) (Fig. 1C and D). These phenotypes could be rescued by expression of *PPC2* (Fig. 1E and F)). Moreover, there were no significant differences in the germination rates between *ppc2* and Col-0 at both CO_2_ conditions (Fig. S1), suggesting the phenotypes of *ppc2* mutant occur in seedling development stage. Compared with Col-0, *ppc2* mutant seedlings accumulated less photosynthetic carbohydrates in leaves, such as sucrose, starch and their upstream precursors (Fig. S2), exogenously addition of sucrose completely recovered the growth arrest phenotype of *ppc2* (Fig. 4A). Mutation of *PPC2* greatly decreased malate content and PEPC activity at low CO_2_ conditions (Figs. 2B, Fig. S2B). Furthermore, in these three plant-type PEPCs in Arabidopsis, only *PPC2* was induced by low CO_2_, consistent with the previous study (Li *et al.*, 2014), and mutation of *PPC1* or *PPC3* did not affect plant growth at low CO_2_ conditions (Figs. 1C and 2A).(Li *et al.*, 2014;. All these results suggest PPC2 is the major PEPC expressed in Arabidopsis leaves and is essential for plant growth at low CO_2_.

### PPC2 participates in photorespiration by linking with primary metabolism at photorespiratory low CO_2_ conditions

Recent studies in sunflower have shown that the malate content closely correlates with photorespiration by metabonomics analyses and C_3_ PEPC fixation increased at high photorespiratory condition (low CO_2_ or high O_2_) (Abadie *et al.*, 2017; Abadie and Tcherkez, 2019), indicating non-photosynthetic PEPC may still contribute to photorespiration in C_3_ plant. The studies in Arabidopsis have reported the novel function of C_3_ PEPC in regulating the balance of carbon/nitrogen metabolism (Masumoto *et al.*, 2010; Shi *et al.*, 2015). However, the molecular mechanisms remain unclear. Here, we were the first to clarify the special role of *PPC2* in photorespiration at low CO_2_ conditions by regulating carbon/nitrogen balance.

Firstly, PPC2 may regulate the balance of carbon/nitrogen at photorespiratory low CO_2_ conditions. The metabolites of glycolysis pathway and amino acid levels in *ppc2* seedlings were greatly affected at photorespiratory low CO_2_ conditions (Fig. 6A; Figs. S2B and S4). The photorespiratory intermediates glycine and serine were significantly accumulated in *ppc2* at photorespiratory CO_2_ conditions (Fig.6A and Fig. S4), suggesting PPC2 is involved in photorespiratory metabolism at low CO_2_ conditions. Glutamic acid plays a central signaling and metabolic role in regulating carbon and nitrogen assimilatory balance (Forde and Lea, 2007), which was greatly reduced in *ppc2* mutant at photorespiratory low CO_2_. The increased glutamine further reduced the assimilation of ammonium released by photorespiration at glycine point, thus likely contributed to the accumulation of glycine and later serine at photorespiratory low CO_2_ conditions. The required glutamate for photorespiration is imported from the chloroplast by exchange against malate, the reduced malate level in the *ppc2* mutant is consistent with this (Fig. S2B). Addition of exogenous malate not only greatly rescued the growth arrest of *ppc2* mutant at low CO_2_ treatment (Fig. 4A), but also reduced the high accumulated photorespiratory intermediates such as glycine in the *ppc1/ppc2* double mutant (Shi *et al.*, 2015). These phenomena are in accordance with the previous report that malate dehydrogenase mutants exhibit an alteration in photorespiratory metabolism (Tomaz *et al.*, 2010). Because of *PPC2* mutation, PEP flux into glycolysis pathway was reduced, and PEP flux into shikimate and calvin cycle was increased at low CO_2_ conditions, leading to higher levels of Phe, Tyr and Trp (Fig. 6A, Fig. S4). These results demonstrate PPC2 control the balance of carbon/nitrogen metabolisms at low CO_2_ conditions.

Secondly, at photorespiratory low CO_2_ conditions *PPC2* affects the expression patterns of photorespiratory enzymes related with glycine and serine synthesis and metabolism. In photorespiratory pathway GGATs and SGAT1 transfers -NH_3_^+^ respectively from glutamate and serine into glyoxylate to synthesize glycine; GLDP1 and GLDT1 decarboxylate glycine into CH_2_-THF; and subsequently SHMT1 transfers the C1 moiety to another glycine resulting in serine formation (Peterhansel *et al.*, 2010). Mutation either in *SGAT1* or *SHMT1* leads to accumulation of serine and glycine (Somerville and Ogren, 1980; Kuhn *et al.*, 2013). Furthermore, overexpression of glycine decarboxylase results in lower glycine and serine content (Timm *et al.*, 2015). In agreement, the expressions of *SHMT1*, *SGAT1*, *GLDP1* and *GLDT1* were greatly reduced in *ppc2* at low CO_2_ conditions, which increased the levels of glycine and serine.

Thirdly, the reduced CO_2_ assimilation of *ppc2* mutant at low CO_2_ conditions was caused by the increased photorespiratory lost in plant. In air condition, *ppc2* mutant exhibited declined CO_2_ assimilation rate in the initial part of A-C_i_ curve (50-400 ppm) (Fig. 8A). However, after reducing O_2_ to inhibit the photorespiration, the reduced CO_2_ assimilation of *ppc2* mutant at low CO_2_ conditions was recovered (Fig. 8C). In agreement, the biomass of *ppc2* mutant was decreased. Moreover, our results showed that the reduced carbon assimilation of *ppc2* at low CO_2_ was not due to stomatal status at low CO_2_ conditions, because the stomatal conductance at the steady state and in response to low CO_2_ shift retained normal in *ppc2* as Col-0 (Fig. 3E and F). In this sense, *PPC2* is specifically involved in photorespiration at low CO_2_ conditions (Fig. 8D).

### ABA regulates photorespiration at low CO_2_ conditions through ABI5

ABA induces gene expression and plays a prominent role in the establishment of stress tolerance. However, the relationships of ABA with low CO_2_ stress and simultaneous photorespiration are still limited. In this study, we found that ABA biosynthesis was heavily blocked at low CO_2_ conditions, possibly due to the decrease in carotenoid accumulation (Fig. 3B), which is the precursor for ABA biosynthesis (Ruiz-Sola and Rodriguez-Concepcion, 2012). When *PPC2* was mutated, the ABA synthesis was further reduced (Fig. 4B). Addition of small amount of ABA largely recovered the *ppc2* seedlings growth (Fig. 4C and D), suggesting that low concentration of ABA is required to stimulate photosynthesis and plant growth at low CO_2_ conditions. In addition, it was reported that exogenous ABA induced the photorespiratory rate in Barley by increasing the activity of GOX (glycolate oxidase) (Popova *et al.*, 1987), our data also showed high ABA treatment led to leaf chlorosis in Col-0 (Fig. 4C and D). These results suggest that higher concentration of ABA in turn could reduce photosynthesis at low CO_2_ stress possibly by promoting carbon flux through photorespiratory cycle.

ABI5 is proposed to be a key player in monitoring environmental conditions during seedling growth (Lopez-Molina *et al.*, 2001) and functions as an intermediate in ABA signaling to regulate seed germination and seedling growth. Our results show that *ABI5* plays a key role in seedling growth at photorespiratory low CO_2_ conditions by regulating the expression of photorespiratory pathway enzymes and ABA regulates plant growth at low CO_2_ conditions through modulating *ABI5* expression. *abi5-1* seedlings growing at low CO_2_ had reduced *F*_*v*_/*F*_*m*_ (Fig. 5C and D). A-C_i_ curves of mature *abi5-1* mutant were reduced at low CO_2_ conditions, which could be fully recovered at non-photorespiratory low O_2_ condition (Figs. 8F and G). In *ppc2* plants, ABA synthesis was reduced and *ABI5* was repressed low CO_2_ conditions, ABA treatment greatly rescued the expression of *ABI5* and growth arrest in *ppc2* (Figs. 4D and 5B). Moreover, the expression levels of photorespiratory enzymes related to glycine and serine synthesis and metabolism were reduced in *ppc2* and *abi5* seedlings at low CO_2_ conditions (Fig. 7A and C). Our further experiments show that ABI5 may regulate *SGAT1* and *GLDT1* by directly binding to their promoters and regulates *SHMT1* indirectly (Fig. 7E). *ABI5* expression rescued their expressions (Fig. 7C) and reduced dry-weight biomass of *ppc2* adult plants (Fig. 8H and I).

In summary, our study reveals that *PPC2* is essential for plant growth at photorespiratory low CO_2_ conditions. It plays important role in carbon assimilation at low CO_2_ conditions via regulating carbon and nitrogen metabolisms during photorespiration. We also identified the important role of ABA in photorespiration and the novel function of *ABI5* in photorespiratory low CO_2_ by regulating transcription of photorespiratory enzyme genes. Our works demonstrate the key role of C_3_ PEPCs in photorespiration at low CO_2_ conditions and may offer clues for future studies to understand the mechanism of C_3_ PEPCs in regulation of photosynthesis and for potential application in crop improvement against photorespiration.

## Supplemental Data

Fig. S1. Seeds germination of WT and *ppc* mutants.

Fig. S2. Photosynthetic carbohydrates were reduced in *ppc2* mutant seedlings at low CO_2_ conditions.

Fig. S3. RT-PCR analyses of *ABI5* expression level in WT, *ppc2* and *ABI5* expressing *ppc2* plants.

Fig. S4. Amino acid content in WT and *ppc2* mutant seedlings.

Fig. S5. Predicted ABRE cis-elements in the promoter regions of photorespiratory enzyme genes.

Table S1. List of primers used in this study.

## Acknowledgement

This work was supported by grants from the National Key Research and Development Program of China (2016YFD0100604), the National Natural Science Foundation of China (31771552) and Fundamental Research Funds for the Central Universities (2662017PY034). The authors are grateful to Dr. Mingqiu Dai (Huazhong Agricultural University) for providing the *abi5-1* mutant seeds. The authors declare that they have no competing interests.

## References

Abadie C, Lothier J, Boex-Fontvieille E, Carroll A, Tcherkez G. 2017. Direct assessment of the metabolic origin of carbon atoms in glutamate from illuminated leaves using (13) C-NMR. New Phytol 216, 1079–1089.

Abadie C, Tcherkez G. 2019. In vivo phosphoenolpyruvate carboxylase activity is controlled by CO_2_ and O_2_ mole fractions and represents a major flux at high photorespiration rates. New Phytol doi: 10.1111/nph.15500.

Baker NR. 2008. Chlorophyll fluorescence: a probe of photosynthesis in vivo. Annu Rev Plant Biol 59, 89–113.

Chollet R, Vidal J, O’Leary MH. 1996. PHOSPHOENOLPYRUVATE CARBOXYLASE: A Ubiquitous, Highly Regulated Enzyme in Plants. Annu Rev Plant Physiol Plant Mol Biol 47, 273–298.

Earley KW, Haag JR, Pontes O, Opper K, Juehne T, Song K, Pikaard CS. 2006. Gateway-compatible vectors for plant functional genomics and proteomics. Plant J 45, 616–629.

Feria AB, Bosch N, Sanchez A, Nieto-Ingelmo AI, de la Osa C, Echevarria C, Garcia-Maurino S, Monreal JA. 2016. Phosphoenolpyruvate carboxylase (PEPC) and PEPC-kinase (PEPC-k) isoenzymes in Arabidopsis thaliana: role in control and abiotic stress conditions. Planta 244, 901–913.

Finkelstein RR, Gampala SSL, Rock CD. 2002. Abscisic acid signaling in seeds and seedlings. Plant Cell 14, S15–S45.

Finkelstein RR, Lynch TJ. 2000. The Arabidopsis abscisic acid response gene *ABI5* encodes a basic leucine zipper transcription factor. Plant Cell 12, 599–609.

Finkelstein RR, Wang ML, Lynch TJ, Rao S, Goodman HM. 1998. The Arabidopsis abscisic acid response locus ABI4 encodes an APETALA 2 domain protein. Plant Cell 10, 1043–1054.

Forde BG, Lea PJ. 2007. Glutamate in plants: metabolism, regulation, and signalling. J Exp Bot 58, 2339–2358.

Gehlen J, Panstruga R, Smets H, Merkelbach S, Kleines M, Porsch P, Fladung M, Becker I, Rademacher T, Hausler RE, et al. 1996. Effects of altered phosphoenolpyruvate carboxylase activities on transgenic C_3_ plant Solanum tuberosum. Plant Mol Biol 32, 831–848.

Gerhart LM, Ward JK. 2010. Plant responses to low CO_2_ of the past. New Phytol 188, 674–695.

Giraudat J, Hauge BM, Valon C, Smalle J, Parcy F, Goodman HM. 1992. Isolation of the Arabidopsis ABI3 gene by positional cloning. Plant Cell 4, 1251–1261.

Hellens RP, Allan AC, Friel EN, Bolitho K, Grafton K, Templeton MD, Karunairetnam S, Gleave AP, Laing WA. 2005. Transient expression vectors for functional genomics, quantification of promoter activity and RNA silencing in plants. Plant Methods 1, 13.

Kiirats O, Lea PJ, Franceschi VR, Edwards GE. 2002. Bundle sheath diffusive resistance to CO_2_ and effectiveness of C_4_ photosynthesis and refixation of photorespired CO_2_ in a C_4_ cycle mutant and wild-type Amaranthus edulis. Plant Physiol 130, 964–976.

Kowalski S, Kopuncova M, Ciesarova Z, Kukurova K. 2017. Free amino acids profile of Polish and Slovak honeys based on LC-MS/MS method without the prior derivatisation. J Food Sci Technol 54, 3716–3723.

Kuhn A, Engqvist MK, Jansen EE, Weber AP, Jakobs C, Maurino VG. 2013. D-2-hydroxyglutarate metabolism is linked to photorespiration in the shm1-1 mutant. Plant Biol (Stuttg) 15, 776–784.

Li Y, Xu J, Haq NU, Zhang H, Zhu XG. 2014. Was low CO_2_ a driving force of C_4_ evolution: Arabidopsis responses to long-term low CO_2_ stress. J Exp Bot 65, 3657–3667.

Liu H, Li X, Xiao J, Wang S. 2012. A convenient method for simultaneous quantification of multiple phytohormones and metabolites: application in study of rice-bacterium interaction. Plant Methods 8, 2.

Lopez-Molina L, Mongrand S, Chua NH. 2001. A postgermination developmental arrest checkpoint is mediated by abscisic acid and requires the ABI5 transcription factor in Arabidopsis. Proc Natl Acad Sci USA 98, 4782–4787.

Ma F, Jazmin LJ, Young JD, Allen DK. 2014. Isotopically nonstationary 13C flux analysis of changes in Arabidopsis thaliana leaf metabolism due to high light acclimation. Proc Natl Acad Sci USA 111, 16967–16972.

Masumoto C, Miyazawa S, Ohkawa H, Fukuda T, Taniguchi Y, Murayama S, Kusano M, Saito K, Fukayama H, Miyao M. 2010. Phosphoenolpyruvate carboxylase intrinsically located in the chloroplast of rice plays a crucial role in ammonium assimilation. Proc Natl Acad Sci USA 107, 5226–5231.

Mouillon JM, Aubert S, Bourguignon J, Gout E, Douce R, Rebeille F. 1999. Glycine and serine catabolism in non-photosynthetic higher plant cells: their role in C1 metabolism. Plant J 20, 197–205.

Novitskaya L, Trevanion SJ, Driscoll S, Foyer CH, Noctor G. 2002. How does photorespiration modulate leaf amino acid contents? A dual approach through modelling and metabolite analysis. Plant Cell Environ 25, 821–835.

Peterhansel C, Horst I, Niessen M, Blume C, Kebeish R, Kürkcüoglu S, Kreuzaler F. 2010. Photorespiration. The Arabidopsis Book 8, e0130–e0130.

Popova LP, Tsonev TD, Vaklinova SG. 1987. A possible role for abscisic Acid in regulation of photosynthetic and photorespiratory carbon metabolism in barley leaves. Plant Physiol 83, 820–824.

Rachmilevitch S, Cousins AB, Bloom AJ. 2004. Nitrate assimilation in plant shoots depends on photorespiration. Proc Natl Acad Sci USA 101, 11506–11510.

Ruiz-Sola MA, Rodriguez-Concepcion M. 2012. Carotenoid biosynthesis in Arabidopsis: a colorful pathway. The Arabidopsis Book 10, e0158.

Saibo NJ, Lourenco T, Oliveira MM. 2009. Transcription factors and regulation of photosynthetic and related metabolism under environmental stresses. Ann Bot 103, 609–623.

Sakuraba Y, Jeong J, Kang MY, Kim J, Paek NC, Choi G. 2014. Phytochrome-interacting transcription factors PIF4 and PIF5 induce leaf senescence in Arabidopsis. Nat Commun 5, 4636.

Seemann JR, Sharkey TD. 1987. The Effect of Abscisic Acid and Other Inhibitors on Photosynthetic Capacity and the Biochemistry of CO(2) Assimilation. Plant Physiol 84, 696–700.

Shi J, Yi K, Liu Y, Xie L, Zhou Z, Chen Y, Hu Z, Zheng T, Liu R, Chen Y, et al. 2015. Phosphoenolpyruvate Carboxylase in Arabidopsis Leaves Plays a Crucial Role in Carbon and Nitrogen Metabolism. Plant Physiol 167, 671–681.

Signora L, De Smet I, Foyer CH, Zhang H. 2001. ABA plays a central role in mediating the regulatory effects of nitrate on root branching in Arabidopsis. Plant J 28, 655–662.

Somerville CR, Ogren WL. 1980. Photorespiration mutants of Arabidopsis thaliana deficient in serine-glyoxylate aminotransferase activity. Proc Natl Acad Sci USA 77, 2684–2687.

Suzuki M, Wang HH, McCarty DR. 2007. Repression of the LEAFY COTYLEDON 1/B3 regulatory network in plant embryo development by VP1/ABSCISIC ACID INSENSITIVE 3-LIKE B3 genes. Plant Physiol 143, 902–911.

Taylor SH, Aspinwall MJ, Blackman CJ, Choat B, Tissue DT, Ghannoum O. 2018. CO_2_ availability influences hydraulic function of C_3_ and C_4_ grass leaves. J Exp Bot 69, 2731–2741.

Tcherkez G, Limami AM. 2019. Net photosynthetic CO_2_ assimilation: more than just CO_2_ and O_2_ reduction cycles. New Phytol 223, 520–529.

Timm S, Wittmiss M, Gamlien S, Ewald R, Florian A, Frank M, Wirtz M, Hell R, Fernie AR, Bauwe H. 2015. Mitochondrial Dihydrolipoyl Dehydrogenase Activity Shapes Photosynthesis and Photorespiration of Arabidopsis thaliana. Plant Cell 27, 1968–1984.

Tomaz T, Bagard M, Pracharoenwattana I, Linden P, Lee CP, Carroll AJ, Stroher E, Smith SM, Gardestrom P, Millar AH. 2010. Mitochondrial malate dehydrogenase lowers leaf respiration and alters photorespiration and plant growth in Arabidopsis. Plant Physiol 154, 1143–1157.

Vidal J, Chollet R. 1997. Regulatory phosphorylation of C_4_ PEP carboxylase. Trends Plant Sci 2, 230–237.

von Caemmerer S. 2013. Steady-state models of photosynthesis. Plant Cell Environ 36, 1617–1630.

Yamaguchi-Shinozaki K, Shinozaki K. 2006. Transcriptional regulatory networks in cellular responses and tolerance to dehydration and cold stresses. Annu Rev Plant Biol 57, 781–803.

Yang Y, Yu X, Song L, An C. 2011. ABI4 activates DGAT1 expression in Arabidopsis seedlings during nitrogen deficiency. Plant Physiol 156, 873–883.

Yoo SD, Cho YH, Sheen J. 2007. Arabidopsis mesophyll protoplasts: a versatile cell system for transient gene expression analysis. Nat Protoc 2, 1565–1572.

